# Relative efficiencies of simple and complex substitution models in estimating divergence times in phylogenomics

**DOI:** 10.1101/2020.02.14.949339

**Authors:** Qiqing Tao, Jose Barba-Montoya, Louise A. Huuki, Mary Kathleen Durnan, Sudhir Kumar

**Author notes:** Correspondence to: Sudhir Kumar Temple University, Philadelphia, PA 19122, USA.

## Abstract

The conventional wisdom in molecular evolution is to apply parameter-rich models of nucleotide and amino acid substitutions for estimating divergence times. However, the actual extent of the difference between time estimates produced by highly complex models compared to those from simple models is yet to be quantified for contemporary datasets that frequently contain sequences from many species and genes. In a reanalysis of many large multispecies alignments from diverse groups of taxa using the same tree topologies and calibrations, we found that the use of the simplest models can produce divergence time estimates and credibility intervals similar to those obtained from the complex models applied in the original studies. This result is surprising because the use of simple models underestimates sequence divergence for all the datasets analyzed. We find three fundamental reasons for the observed robustness of time estimates to model complexity in many practical datasets. First, the estimates of branch lengths and node-to-tip distances under the simplest model show an approximately linear relationship with those produced by using the most complex models applied, especially for datasets with many sequences. Second, relaxed clock methods automatically adjust rates on branches that experience considerable underestimation of sequence divergences, resulting in time estimates that are similar to those from complex models. And, third, the inclusion of even a few good calibrations in an analysis can reduce the difference in time estimates from simple and complex models. The robustness of time estimates to models complexity in these empirical data analyses is encouraging, because all phylogenomics studies use statistical models that are oversimplified descriptions of actual evolutionary substitution processes.

## Introduction

Models of nucleotide and amino acid substitution are of fundamental importance in molecular phylogenetic analyses (Nei and Kumar 2000; Yang 2006; Arenas 2015). Many sophisticated substitution models have been developed, and the complexity of models developed for use in phylogenomic studies continues to increase (Arenas 2015; Abadi et al. 2019). Indeed, complex models can provide a more complete description of nucleotide and amino acid substitution processes that involve transition/transversion rate differences, biased base compositions, inequality of evolutionary rates among sites, and substitution pattern heterogeneity among genomic regions and sequence partitions (Sumner et al. 2012; Arenas 2015). Because the difference between the estimated and actual numbers of substitutions grows quickly and non-linearly over time (Nei and Kumar 2000; Yang 2006), researchers often select the most complex model available to improve the accuracy of divergence time estimates (Arbogast et al. 2002; Sumner et al. 2012; Arenas 2015; Abadi et al. 2019).

Do the time estimates from the most complex models differ significantly from those produced using a relatively simple model when analyzing large datasets that contain many species, genes, and calibrations? The answer to this question is of high practical significance. If it is affirmative, time estimates may be vulnerable to both incorrect model specification and the overall limitations of current substitution models. On the other hand, if time estimates are generally similar for simple and complex substitution models, then currently available models could be deemed sufficient for molecular dating, because even the most complex model is a simplification of actual evolutionary processes that are influenced by differences in regional mutation patterns and spatial and temporal selective pressures (Arenas 2015; Abadi et al. 2019).

While no studies have directly examined the impact of model complexity on time estimation in phylogenomic investigations, there have been reports that simple substitution models often perform similarly to or slightly worse than complex substitution models in some types of phylogenetic inferences (Tamura et al. 2004; Yoshida and Nei 2016; Dornburg et al. 2018; Spielman and Kosakovsky Pond 2018; Abadi et al. 2019; Spielman 2019). However, in molecular dating, it is intuitively assumed that underestimation of sequence divergence caused by the use of simple models will result in significantly distorted time estimates. Here, we used diverse large datasets to test the conventional wisdom that the use of simple models will result in poor estimates of divergence time (**Table 1**). To detect potential benefits offered by the use of complex models, we compared Bayesian estimates of divergence times obtained when using extremely simple models with those obtained when using complex models that contained many biological attributes and parameters (**Table 1**).

**Table 1.** Detailed information about the empirical data analyzed.

| Taxonomic group | Data type<sup>a</sup> | Sequence count | Sequence length<sup>b</sup> | Tree depth<sup>c</sup> | Rate<sup>d</sup> | Substitution model<sup>e</sup> | Partition count | Rate model<sup>f</sup> | Calibration count | $\Delta \ln L^g$ | Reference |
| --- | --- | --- | --- | --- | --- | --- | --- | --- | --- | --- | --- |
| Mammals (A) | M | 274 | 7,370 | 185 | 2.28 | HKY + $\Gamma$ | 1 | ABR | 36 | 96320.5 | dos Reis et al. (2012) |
| Mammals (B) | A | 162 | 11,010 | 187 | 1.53 | JTT + $\Gamma$ | 26 | ARB | 64 | 110854.0 | Meredith et al. (2011) |
| Birds | N | 51 | 722,202 | 102 | 0.21 | HKY + $\Gamma$ | NA | ABR | 18 | 3794.2 | Jarvis et al. (2014) |
| Fishes | N | 118 | 85,363 | 140 | 0.51 | HKY + $\Gamma$ | 4 | IBR | 13 | 13048.1 | Alfaro et al. (2018) |
| Metazoans | A | 54 | 38,577 | 757 | 0.41 | LG + $\Gamma$ + F | 1 | IBR | 33 | 68405.1 | dos Reis et al. (2015) |
| Spiders | A | 43 | 55,447 | 561 | 0.54 | WAG + $\Gamma$ | 1 | IBR | 8 | 37088.5 | Bond et al. (2014) |
| Plants | N | 103 | 856,439 | 798 | 0.87 | GTR + $\Gamma$ | 1 | IBR | 37 | 14757.9 | Morris et al. (2018) |
| Eukaryotes & Prokaryotes | A | 102 | 9,874 | 4511 | 0.29 | LG + $\Gamma$ | 29 | IBR | 11 | 183666.0 | Betts et al. (2018) |

<sup>a</sup>N = nuclear DNA; M = mitochondrial DNA; A = amino acid.
<sup>b</sup>Sequence length is the number of sites in the alignment used in the original study. We randomly selected 10,000 sites in our analyses if sequences are longer than 10,000 sites. For the 'Plants' dataset, we used an alignment of 2,217 sites.
<sup>c</sup>Tree depth is the root age obtained from the original published study. Times are in millions of years.
<sup>d</sup>The evolutionary rate is calculated by dividing the sum of the maximum likelihood (ML) branch lengths obtained using the original complex model over the sum of published times elapsed. The unit is substitutions per site per billion years.
<sup>e</sup>The substitution model used for the majority of partitions in the original study for estimating divergence times.
<sup>f</sup>The branch rate model used in the original study for estimating divergence times. ABR: autocorrelated branch rate model; IBR: independent branch rate model.
<sup>g</sup>Difference between the log-likelihoods of the original complex model and the simple model. A positive value means that the complex model provides a better fit to the data.

We focus on models of the General Time Reversible (GTR) class because they are employed in all current empirical dating analyses. Although many new substitution models have been proposed to relax the assumptions of stationarity, reversibility, and homogeneity of base substitution patterns, as well as to incorporate complex structural constraints and epistatic fitness landscapes, these advanced models are not yet available in popular phylogenetic software packages due to their implicit complexity and onerous computational requirements (Jayaswal et al. 2014; Arenas 2015; Arenas et al. 2015; Usmanova et al. 2015). Therefore, in our reanalysis, we used the models selected as the best-fit models in the source phylogenomic studies as ‘complex models’ (**Table 1**). The ‘simple models’ used were the Jukes-Cantor (JC) model for nucleotide substitutions (Jukes and Cantor 1969) and the Poisson model for amino acid substitutions (Nei and Kumar 2000). Both assume that all substitution types are equally likely at a given site, an assumption that is always violated in reality. To ensure the most powerful contrast, we applied these simplest models without partitioning the sequence alignment by genes, genomic features (e.g., codon positions), or sets of positions with similar substitution patterns (see ***Materials and Methods***).

We also used the same tree topology as in the source publications in our comparisons, because it is a common practice in phylogenomic studies to use the phylogeny reliably inferred using a sophisticated tree-building method as a fixed topology for molecular dating (e.g., Li et al. 2019; Oliveros et al. 2019). Also, the use of the same published tree topologies ensured consistent placement of calibrations and eliminated the confounding effects of using alternative phylogenies and calibrations.

## Results

### A plant dataset analysis

We first present results from reanalysis of a large sequence alignment containing 103 plant species (Morris et al. 2018) (‘Plants,’ **Table 1**). Pairwise sequence distances ranged from 0.01 – 3.18 (median = 0.83) nucleotide substitutions per site (**Fig. 1a**). The GTR model with rate variability among sites (+Γ) was used in the original analyses, and is presented here as the complex model (Morris et al. 2018). For comparison, we analyzed this dataset using the JC model as our simple model. The difference in the maximum likelihood (ML) values for the GTR+ Γ and JC models was very large (Δ*ln*L = 14757.9) and highly significant (*P* < 10^−16^). Conventionally, the JC model would be a poor choice for molecular dating analysis. Indeed, the use of the JC model led to the underestimation of pairwise evolutionary distances by as much as 73%, and there was severe substitution saturation resulting in a classic curvilinear trend (**Fig. 1a,** the gray area).

**Figure 1.**
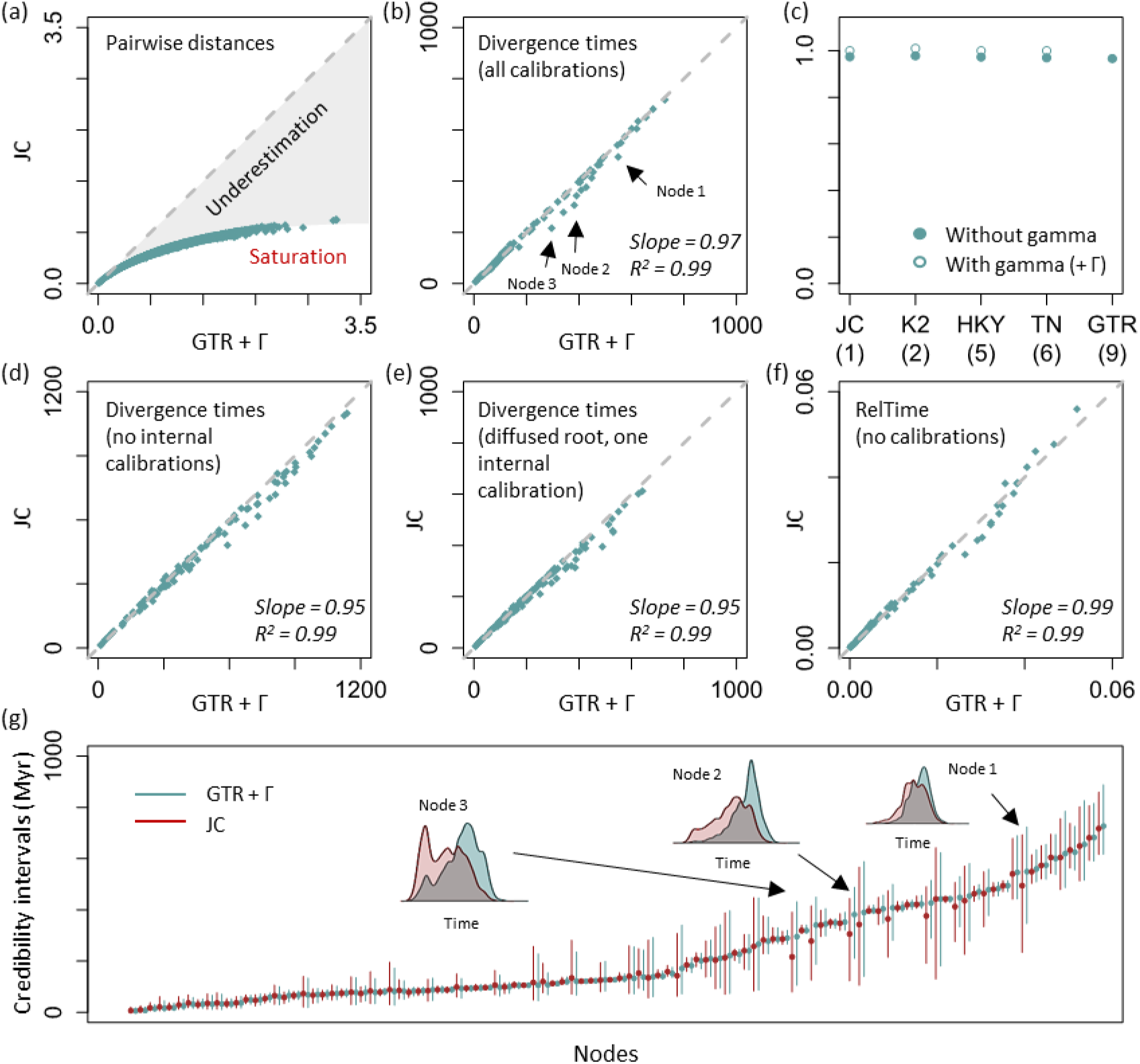
Plant data analyses. (a) Severe underestimation of pairwise distances via the JC model. The gray dashed line represents equality between time estimates and the gray area represents the underestimation resulting from using the JC model. (b) Similar divergence time estimates are produced by using the JC and GTR + Γ models when all calibrations are used in Bayesian analyses. Gray dashed line represents 1:1 line. The slope and coefficient of determination (*R*^2^) for the linear regression through the origin are shown. Arrows mark three nodes that show different time estimates. (c) Relationship between the complexity of models and the slope of divergence times inferred using the GTR + Γ and other models. JC, K2, HKY, TN, and GTR represents Jukes-Cantor, Kimura-2-parameter, Hasegawa-Kishino-Yano, Tamura-Nei, and General Time Reversible models, respectively. The number of model parameters is shown in the parentheses. Circles indicate whether a gamma distribution (+ Γ) for incorporating rate variation across sites is used (open circle) or is not used (closed circle) with the substitution model. The Bayesian method produces similar estimates between the JC and GTR + Γ models when (d) all internal calibrations are excluded, and (e) one internal calibration and a diffused root calibration are used. (f) The RelTime method produces similar divergence times between the JC and GTR + Γ models. Times are normalized to the sum of node ages. (g) Comparison of 95% Bayesian credibility intervals generated under the JC (dark red) and GTR + Γ (cadet blue) models. Dots are point estimates of divergence times. Distributions of posterior time estimates for three nodes pointed in panel b are shown (inset).

Surprisingly, Bayesian estimates of divergence times obtained using the JC model were very similar to those obtained using the GTR + Γ model when the same sequence alignment, topology, and calibrations were used (**Fig. 1b**). This trend was also observed for divergence time estimates obtained via substitution models of intermediate complexity (**Fig. 1c**). The linear regression slope between time estimates under the JC and GTR + Γ models was 0.97, with low dispersion (*R*^2^ = 0.99). Although time estimates generated using the GTR+ Γ and JC models showed high overall similarity, a few node times showed local discrepancies (e.g., arrows in **Fig. 1b**). However, these node times generated using the JC model fell within the 95% credibility intervals (CrIs) obtained using the GTR + Γ model (**Fig. 1g**). CrIs from the JC and GTR + Γ models overlapped for every node (**Fig. 1g**), suggesting that estimates of divergence times and CrIs from the simplest model will be as useful as those obtained via complex substitution models in downstream biological analyses and hypothesis testing.

We hypothesized that the inclusion of 37 calibration points, and their associated probability densities, in the ‘Plants’ dataset constrained the node time estimates and eliminated the bias anticipated to be caused by the use of the simple JC model. So, we compared times obtained using the GTR+ Γ and JC models after eliminating all the internal calibrations but retaining the original root calibration. The linear pattern persisted (slope = 0.95, *R*^2^ = 0.99, **Fig. 1d**). We then tested the possibility that a well-constrained root calibration caused the observed linear relationship of dates from simple and complex models, even though the root calibration was expected to only dictate the overall time span, rather than the patterns of individual node times. So, we reanalyzed the phylogeny, making the probability density distribution of the root calibration diffused and adding a randomly selected internal calibration in the Bayesian analyses (see ***Materials and Methods***). The resulting divergence times from using the simple JC model were still very similar to those from the more complex GTR + Γ model (slope = 0.95, *R*^2^ = 0.99, **Fig. 1e**). Therefore, dates from simple and complex models show good linear relationships in Bayesian analyses with even a few calibrations.

To eliminate the effect of calibrations in mediating the similarity of times obtained using simple and complex models, we estimated divergence times using the RelTime method in which no calibrations and no branch rate model are required. For this dataset, divergence times under the JC and GTR + Γ models were very similar (**Fig. 1f**) and the confidence intervals also showed broad overlap. This result suggested that the specification of a branch rate model and calibrations used in Bayesian analysis for the datasets analyzed are unlikely to explain the high similarity in divergence times between the JC and GTR + Γ model. Overall, substitution model complexity appears to have limited impact on divergence time estimates for the ‘Plants’ dataset.

### The complexity of the substitution model has limited impact on time inference for many datasets

We examined the similarity of times estimated via simple and complex models for many other datasets, which contained small or large numbers of species (43 - 274), had varying evolutionary time depths (102 – 4511 million years, Myrs), evolved with slow or fast rates (0.21 – 2.28 substitution per site per billion years), or employed small or large numbers of calibration points (8 - 64) (**Table 1**). For all these datasets, the use of simple models underestimated pairwise evolutionary distances and showed curvilinear relationships with distances estimated using complex models (**Fig. 2**). The curvilinear relationship was particularly dramatic for more ancient divergences due to substitutional saturation (**Fig. 2a**).

**Figure 2.**
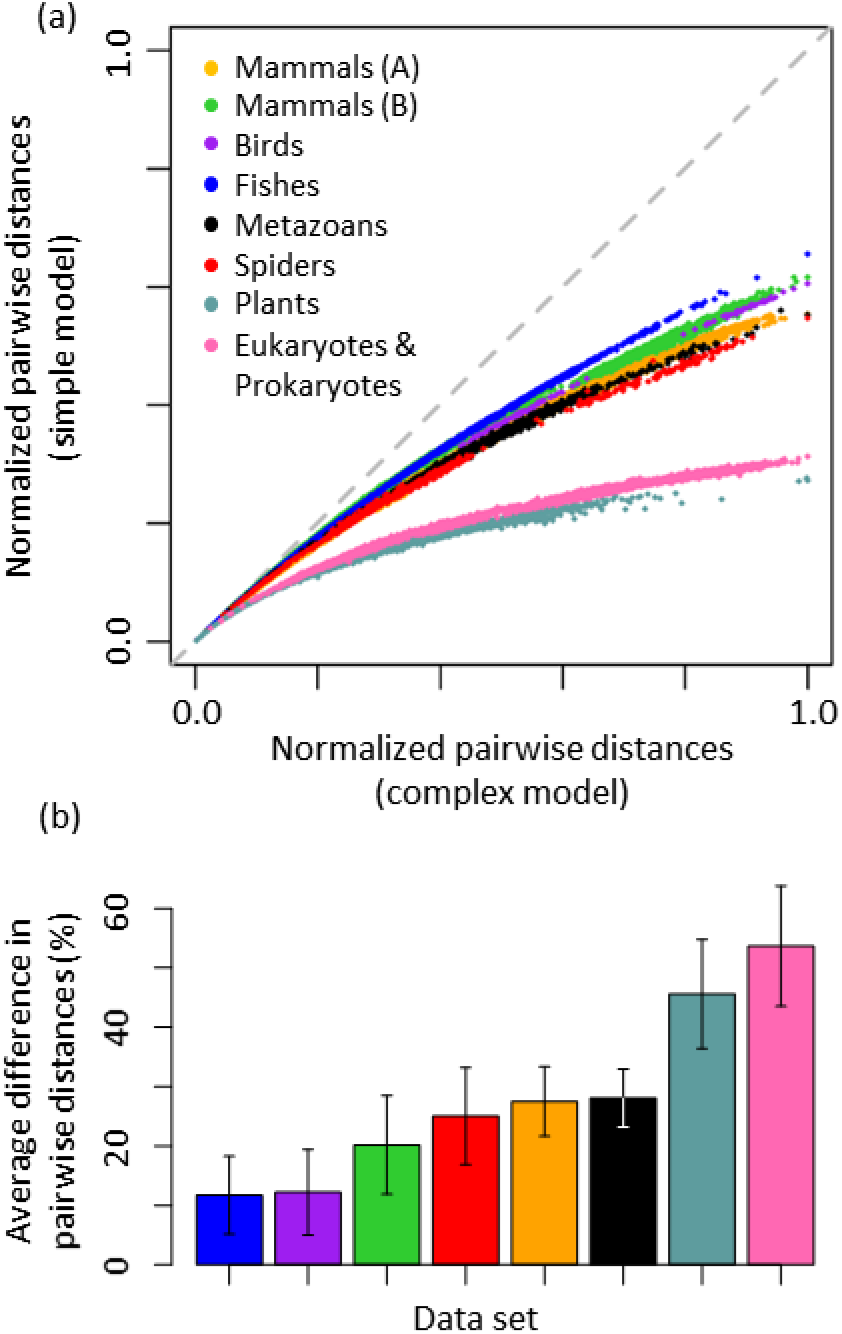
(a) Curvilinear relationships of pairwise distances. Pairwise distances are normalized to the maximum pairwise distance obtained using the complex model for a given empirical dataset to enable comparisons across empirical datasets. The gray dashed line represents equality between distance estimates. (b) Average percent differences between pairwise distances obtained using simple and complex models. The error bar shows one standard deviation. Datasets for ‘Mammals (A)’, ‘Mammals (B)’, ‘Birds’, ‘Fishes’, ‘Metazoans’, ‘Spiders’, ‘Plants’, and ‘Eukaryotes & Prokaryotes’ are derived from dos Reis et al. (2012), Meredith et al. (2011), Jarvis et al. (2014), Alfaro and Holder (2006), dos Reis et al. (2015), Bond et al. (2014), Morris et al. (2018), and Betts et al. (2018).

Despite the curvilinear relationship of pairwise distances estimated using simple and complex models, their divergence time estimates showed strong linear relationships for nuclear nucleotide, mitochondrial nucleotide, and amino acid sequence alignments (**Fig 3a**). The linear regression slopes ranged from 0.92 to 1.01 (**Fig. 3a**). The linear relationships persisted in all analyses even when internal calibrations were removed (only the root calibration was kept); slopes ranged from 0.96 to 1.11 (**Fig. 3b**). The use of a random internal calibration and a diffused root calibration produced a similar range of slopes (0.94 to 1.01) (**Fig. S1**). As with the Plants dataset, RelTime dating analysis also produced excellent linear relationships of node times estimated via simple and complex models (**Fig. S2**).

**Figure 3.**
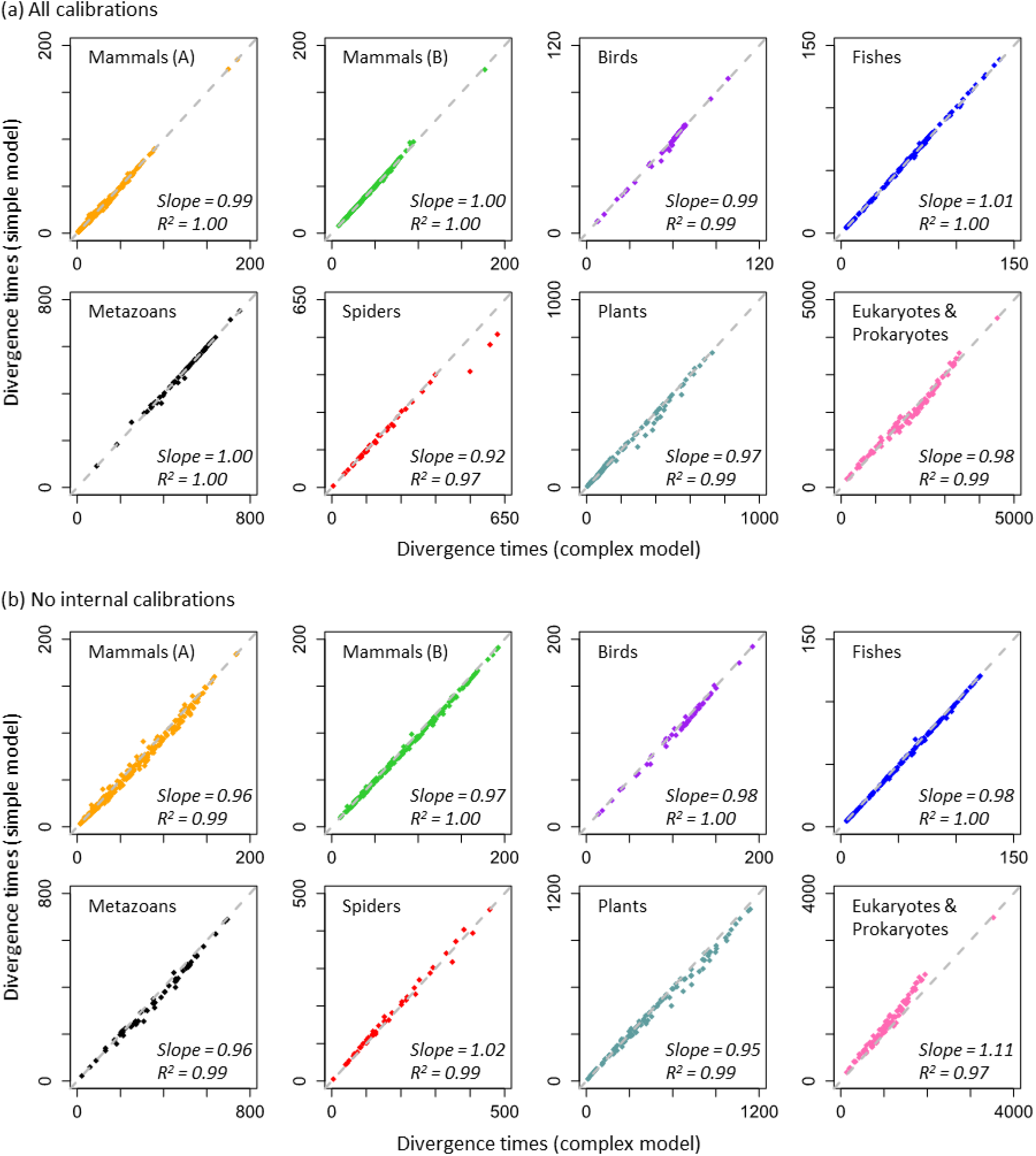
Comparisons of Bayesian divergence times obtained via simple and complex models. Similar divergence time estimates are produced when (a) all calibrations are used, and (b) all internal calibrations are excluded in Bayesian analyses. The time unit is millions of years. The gray dashed line marks equal time estimates. The slope and coefficient of determination (*R*^2^) for the linear regression through the origin are shown. Source publications for datasets are listed in figure 2.

The mean of the relative absolute difference between times estimated by simple and complex models was small (< 6.2%), but some node times deviated considerably from the 1:1 linear trend for these two models. For example, three nodes in the ‘Spiders’ dataset showed 15% - 23% difference between time estimates obtained from simple and complex models (**Fig. 3a**). Interestingly, these differences disappeared (< 1%) when all the internal calibrations were excluded (**Fig. 3b**). In contrast to the patterns observed for the ‘Spiders’ dataset, divergence estimates from the ‘Eukaryotes & Prokaryotes’ dataset showed larger differences between simple and complex models when all the internal calibrations were removed. However, Bayesian CrIs obtained using simple models often contained the point estimates obtained using complex models, and vice versa (**Fig. 4** and **Fig. S3**). CrIs overlapped for more than 97% of the nodes across all the analyses between simple and complex models (**Fig. 4**) and the overlapping region was significant for the majority of the nodes (**Fig. S4**). Therefore, simple and complex models seem to offer similar statistical power for biological hypothesis testing. For example, in the analysis of ‘Mammals (A)’ dataset, divergence estimates from both simple (88.6 – 91.3 Myr) and complex (88 – 90.6 Myr) models rejected the evolutionary model in which the last common ancestor of placental mammals appeared after the Cretaceous-Paleogene (K-Pg) event, consistent with conclusions in some previous studies (Hedges et al. 1996; Kumar and Hedges 1998; dos Reis et al. 2012).

**Figure 4.**
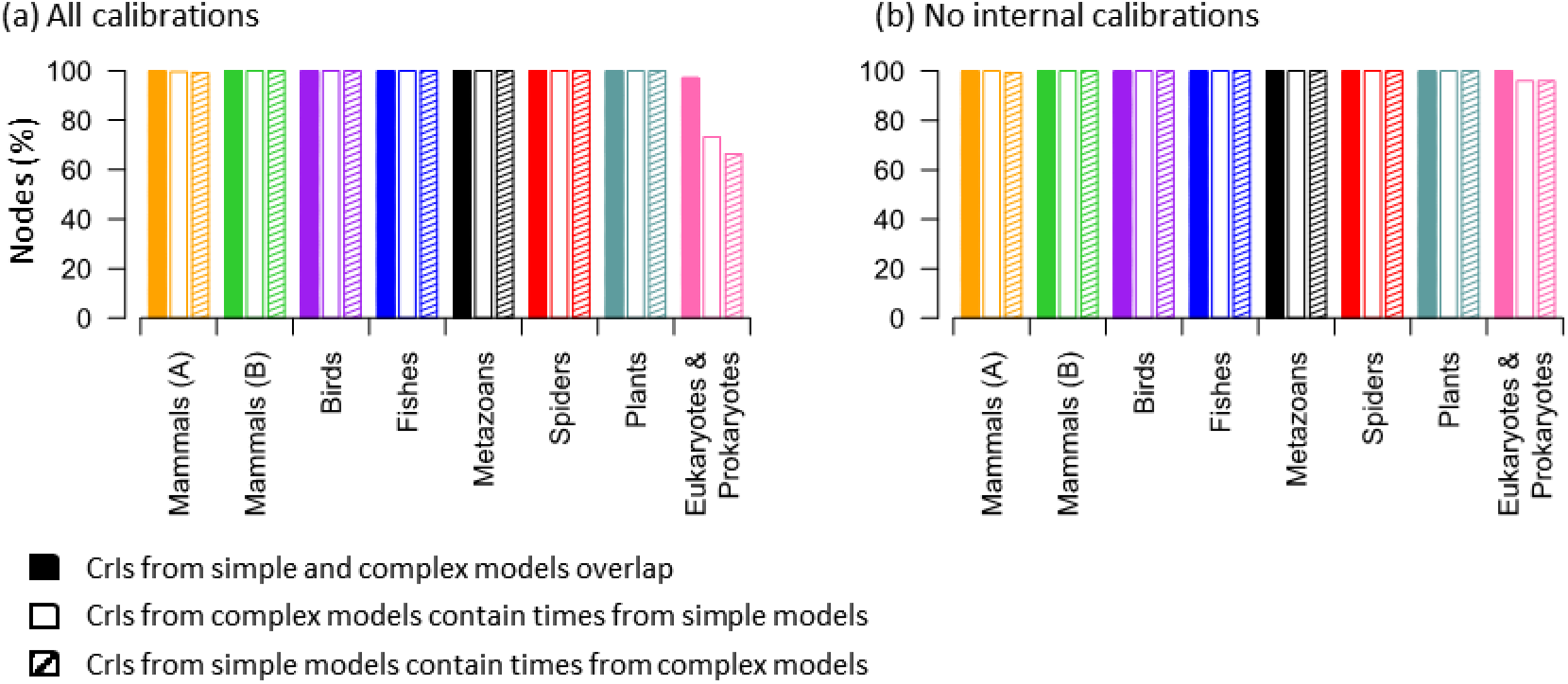
Comparisons of Bayesian credibility intervals (CrIs) inferred by using simple and complex models. Shown are the proportions of node times for which simple and complex models produce overlapping CrIs (solid), CrIs produced by complex models contain point time estimates produced by simple models (open), and CrIs produced by simple models include point time estimates produced by complex models (hatch) when (a) all calibrations are used and (b) no internal calibrations are used. See also figure S3 for more detailed information about panel **a**.

### A fundamental factor enabling robustness of inferred times to model complexity

Elucidation of causal factors mediating the similarity of time estimates produced under simple and complex substitution models is needed to reveal the fundamental basis of our observation and to establish its generality. We hypothesized that the branch length estimates obtained via simple and complex models were linearly related (even though not 1:1) for the datasets analyzed. This linearity would result in similar relative branch lengths, and thus divergence times. This hypothesis arose from our observation that the RelTime method produced similar time estimates under simple and complex models (**Fig. S2**). RelTime is based on a relative rate framework in which the relationship between branch lengths and time estimates is established algebraically (Tamura et al. 2012; Tamura et al. 2018). Divergence times are the ratios of the linear combinations of branch length estimates (and of node-to-tip distances), which allows us to predict that the relative branch lengths between simple and complex models will be similar.

Indeed, linear models (through the origin) described the relationship between branch lengths from simple and complex models for all empirical datasets we examined (**Fig. 5**). This trend is dramatically different from that observed for pairwise distances, where a curvilinear relationship was observed in every case (**Fig. 2**). Linear regression slopes of branch lengths are all less than 1 because simple models underestimate sequence divergences. However, branch lengths were uniformly underestimated via simple models for the empirical datasets tested, resulting in similar relative branch lengths between simple and complex model scenerios (**Fig. 5**) as well as similar relative node-to-tip distances (**Fig. S5**). Because divergence times are a function of the ratios of the linear combinations of branch lengths, and the ratios of node-to-tip distances, they become comparable under simple and complex models.

**Figure 5.**
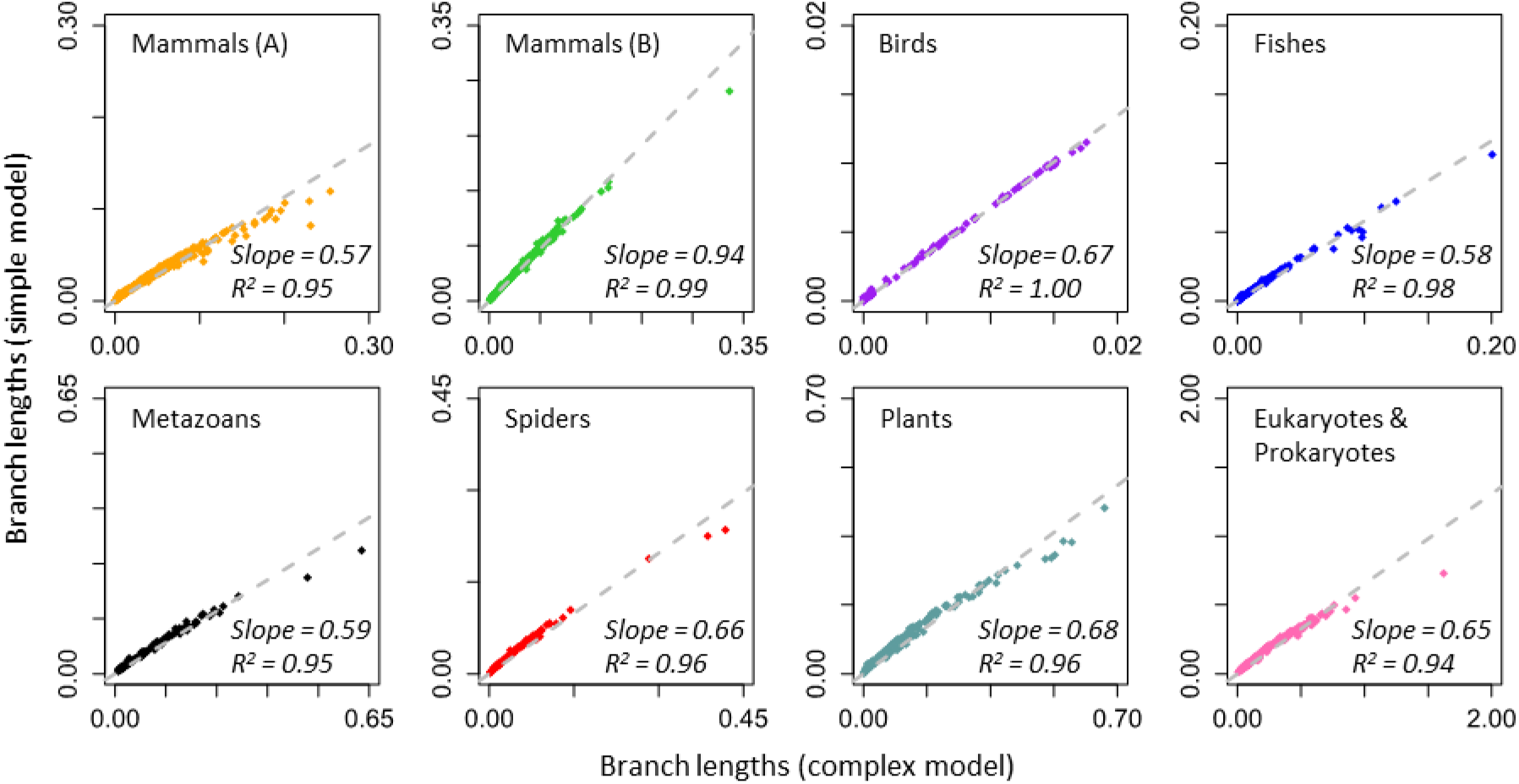
Relationship between ML branch lengths obtained by using simple and complex models. The gray dashed line represents the best-fit linear regression through the origin. The slope and coefficient of determination (*R*^2^) are shown.

Despite the linear patterns described above, we expected a greater magnitude of underestimation for longer branches and deeper sequence divergences when using simple models (see **Fig. 2a**), which could distort time estimates. This effect is indeed observed for the longest branches in most of the datasets analyzed, as they show significant deviation from the linear trend (**Fig. 5**). We confirmed this pattern in a systematic analysis of short, long, and intermediate branch lengths (**Fig. 6a**); and shallow, deep, and intermediate node-to-tip distances (**Fig. 6c**). However, divergence time estimates on deep nodes and branch times on long branches were often not very different between simple and complex models (**Fig. 6b** and **d**). These results suggest that relaxed clock methods automatically adjust evolutionary rates within a phylogeny to produce robust time estimates for the empirical datasets analyzed.

**Figure 6.**
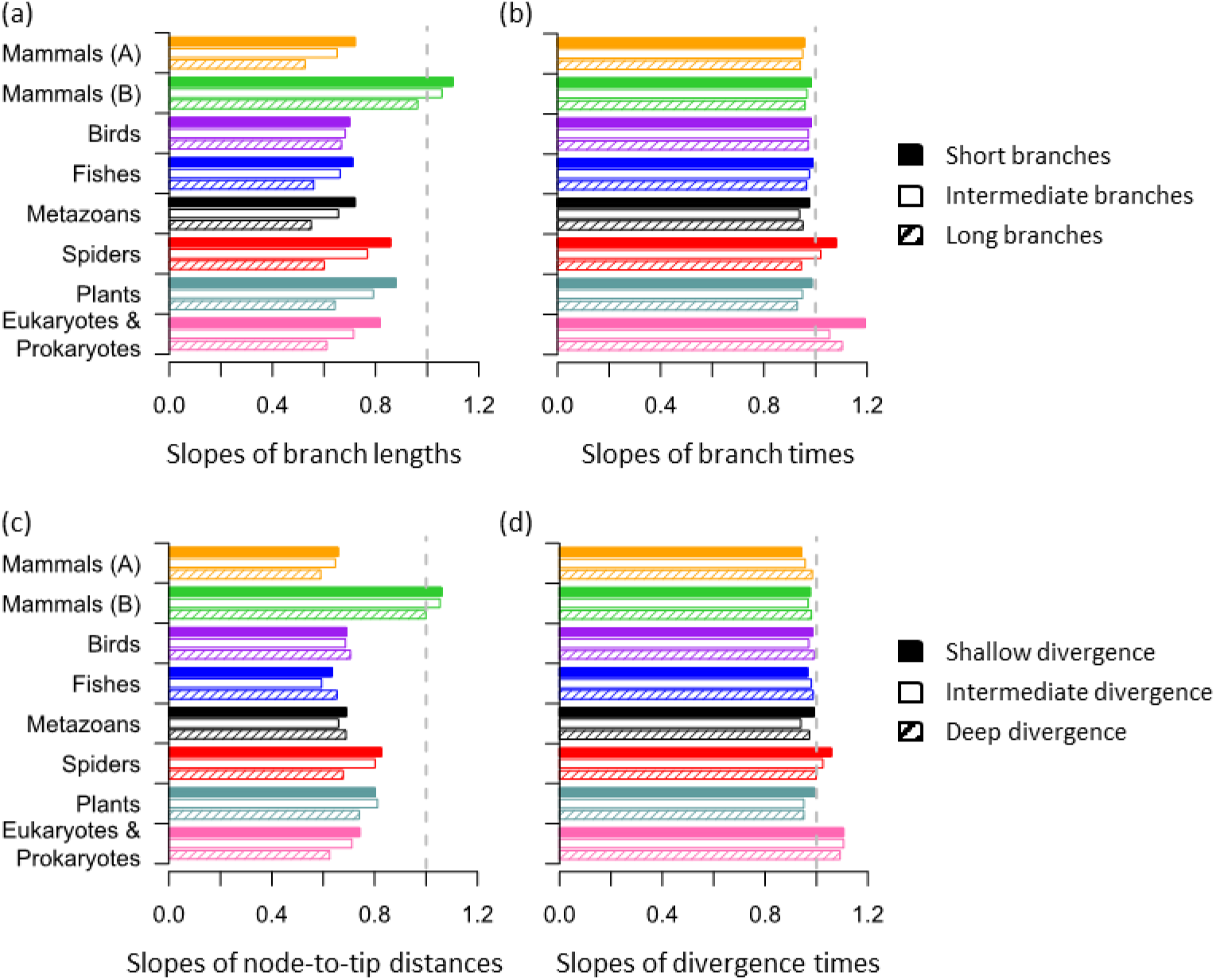
(a) Linear regression slopes of branch length and (b) branch time estimated using simple and complex models for short (solid), intermediate (open), and long (hatch) branches. (c) Linear regression slopes of node-to-tip distance and (d) divergence time estimated using simple and complex models for shallow (solid), intermediate (open), and deep (hatch) locations in the phylogeny. A slope of 1 represents equality between estimates from simple and complex models, which is marked by a gray dashed line. Smaller slope values represent more considerable underestimation when using simple models.

However, this adjustment may not be possible for phylogenies with few species or sparse taxon sampling within some clades, producing isolated long branches and causing time estimates from simple models to differ from complex models. For example, many very long branches may appear in an unbalanced phylogeny (**Fig. S6a**). Interestingly, we found that the slope of times estimated using simple and complex models was close to 1 when the rates were similar to those observed in empirical datasets (1× category in **Fig. S6b** and **c**), which is consistent with results from our empirical analyses. For datasets where evolutionary rates are extremely fast (10×), or sequence divergences are very large (see ***Materials and Methods***), we found that the use of a single, very shallow calibration often produced overly young estimates (blue dots, 2× - 10× in **Fig. S6c**). It is because of a more severe underestimation of branch lengths for long branches when usinga simple model, and the rate adjustment offered by relaxed clock methods is not as effective. However, the use of calibrations at deeper nodes alleviated the discrepancy and produced less biased time estimates for the simple model (green and pink dots, 2× - 10× in **Fig. S6c**). Because reliable alignment of sequences showing large divergences is very challenging (Edwards et al. 1995), researchers will (and should) generally be apprehensive of using highly divergent genes (e.g., 10×).

### Increasing numbers of sequences make time estimates from simple and complex models more similar

We investigated the effect of the number of sequences in a dataset on the similarity of estimated dates from simple and complex models, because a dataset with sparse taxon sampling may show greater discrepancy. We evaluated the linearity of the relationship of branch lengths generated via simple and complex models for data subsets with increasingly larger numbers of sequences subsampled from the ‘Plants’ dataset. We found that when the number of sequences sampled was small (e.g., 10), the linear relationship between branch lengths was weak for some subsets (lower linear coefficient, *R*^2^) and strong for others (higher *R*^2^) (**Fig. 7**). With an increasing number of sequences, the dispersion of *R*^2^ values became smaller, resulting in a more robust linear relationship between branch lengths (**Fig. 7**). In the empirical datasets analyzed, the dispersion of *R*^2^ became very small for datasets that contain as few as 40 sequences. Therefore, a stronger linear relationship of branch length estimates between simple and complex models exists for datasets with many species, resulting in similar divergence time estimates. Our results may explain the inconsistent patterns reported in previous studies (Yang 1996; Schenk and Hufford 2010), which analyzed relatively small datasets (5 - 25 sequences).

**Figure 7.**
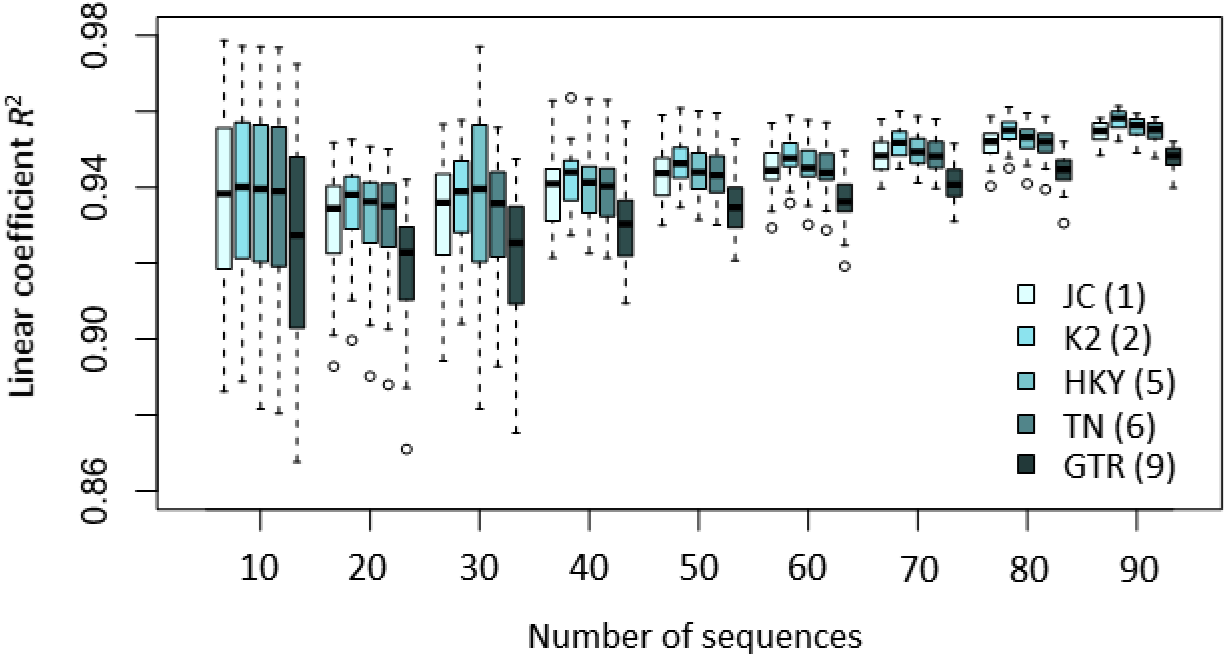
Relationships between the number of sequences and the dispersion around the linear trends of branch lengths from simple and complex models. Boxes show the variation of the coefficient of determination of the linear regression (through the origin, *R*^2^) between branch lengths obtained using the GTR + Γ and the corresponding models (JC, K2, HKY, TN, and GTR) based on an analysis of 20 replicate datasets. A narrower box indicates a more stable linear relationship of branch lengths. Model abbreviations are as those in figure 1. The number of model parameters is shown in the parentheses.

### Effects of irreversibility and non-stationarity of substitution patterns

As mentioned earlier, our primary focus is on complex models used in practical analyses, which are from the GTR class. However, it is important to consider whether violation of the assumption of model stationarity and time-reversibility has biased time estimates significantly. Because non-GTR substitution models are not available for use in the Bayesian dating software, we used the RelTime method to infer divergence times directly for phylogenies in which branch lengths were estimated under an unrestricted model (in which the time-reversibility in substitution models is not assumed) and a model in which the stationarity of the substitution patterns is not assumed (see ***Materials and Methods***). We found that the JC model and non-GTR models produced similar divergence time estimates and confidence intervals (**Fig. 8a** and **b**). It is because branch length estimates obtained using the JC and non-GTR models showed strong linear relationships (**Fig. 8c** and **d**). Simple and complex models generated comparable time estimates and, hence, may provide equivalent statistical power for hypothesis testing within the range of evolutionary conditions observed in empirical datasets analyzed for this study.

**Figure 8.**
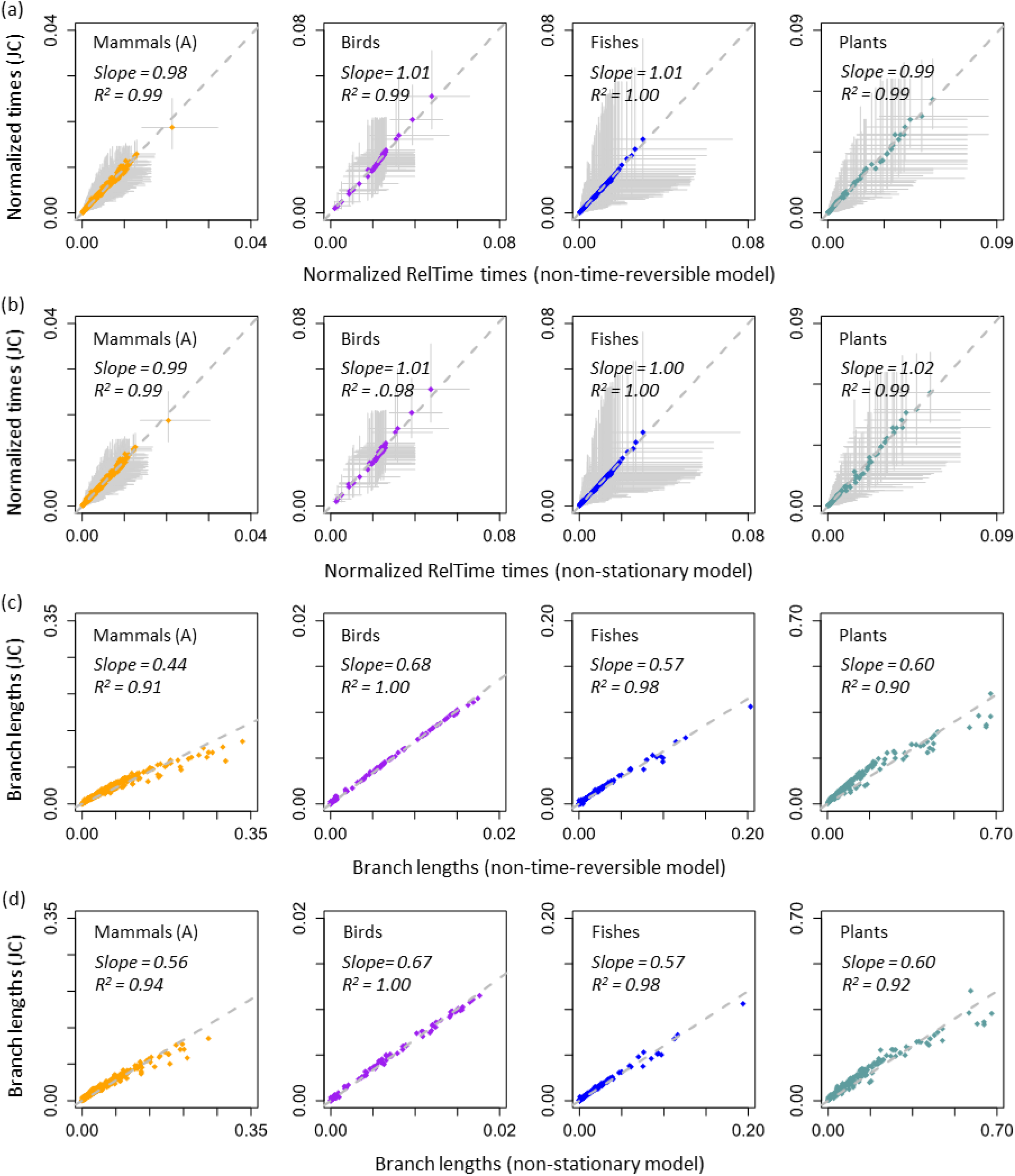
Relationships of RelTime divergence times estimated with branch lengths obtained using the JC model and models that are (a) non-time-reversible and (b) non-stationary for all nucleotide datasets. Gray solid lines represent 95% confidence intervals. The sum of node ages was used to normalize divergence times and confidence intervals. Relationships of branch lengths obtained using the JC model and models that are (c) non-time-reversible and (d) non-stationary. The gray dashed line represents the best-fit linear regression through the origin. The slope and coefficient of determination (*R*^2^) for the linear regression are shown.

## Discussion

Divergence times estimated using simple and complex substitution models were remarkably similar across a range of phylogenomic datasets, a pattern that we observed for both Bayesian and RelTime methods. More surprisingly, similar estimates were observed even when a small number of calibrations (e.g., only the root calibration) was used in Bayesian and non-Bayesian analyses. We found that three fundamental reasons can explain the observed robustness of time estimates to model complexity in many phylogenomic datasets. First, the estimates of branch lengths and node-to-tip distances under simple and complex models show strong linear relationships, especially for datasets with many sequences. Second, relaxed clock methods are able to automatically adjust evolutionary rates on branches that experience underestimation of sequence divergence, resulting in time estimates that are similar to those from complex models. Third, the use of calibrations (especially deep calibrations) further narrows differences in time estimates from simple and complex models, as calibrations often offer strong constraints on node ages.

Although divergence time estimates derived via simple and complex models show remarkable overall similarity, use of different substitution models may produce very different point estimates for some nodes (e.g., >20% difference for node 3 in **Fig. 1g**). It is mainly because the distributions of posterior times under simple and complex models differ for those nodes. Even for phylogenomic datasets, divergence time estiamtes are generally associated with large estimation errors due to variance of branch length estimates, the degree of evolutionary rate heterogeneity, and uncertainty related to clock calibration (Zhu et al. 2015; Tao et al. 2020).Therefore, CrIs, which represent the uncertainty suttouding divergence time estiamtes, are more useful than point estimates in biological hypothesis testing (Warnock et al. 2017). CrIs produced by simple and complex models largely overlapped (**Fig. 4** and **Fig. S4**), and the widths of CrIs around time estimates were also very similar when the same set of calibrations were used (**Fig. 1g** and **Fig. S3**).

Even though we found that simple models produced results comparable to those from complex models across many empirical datasets, we anticipate that there will be situations in which complex models are best suited for molecular dating. This includes the analysis of datasets in which the number of sequences is small, substitution patterns have shifted substantially in some groups, sequences divergences are large, or taxon sampling in some clades is so sparse as to create many long branches. Complex models are also required when using an external mutation or substitution rate to set evolutionary distances to time because actual branch lengths are best estimated via a complex model. Therefore, complex models are needed if one is interested in estimating not only divergence times but also absolute and relative evolutionary rates. Complex models may also be required when inferring the phylogeny and divergence times jointly, although use of a predetermined topology is a common practice in phylogenomic studies (e.g., Li et al. 2019; Oliveros et al. 2019). In the future, we plan to investigate the impact of substitution model complexity on the joint inference of phylogeny and times.

A majority of phylogenomic analyses make a simplifying assumption that the same substitution model applies across all the sites in a concatenated dataset or within each data partition. However, evolutionary dynamics and processes differ regionally (and even positionally) in mutation patterns, sequence context, and selective pressures (Yang et al. 1994; Yang and Swanson 2002; Shapiro et al. 2005; Kosakovsky Pond et al. 2008; Bordner and Mittelmann 2013; Jayaswal et al. 2014; Arenas 2015). Therefore, one may imagine that even a seemingly complex substitution model (e.g., GTR + Γ) is relatively simple when compared to the actual reality. The observation that the simplest models produce time estimates similar to those obtained using much more complex models may be used to suggest that the current model modelxity is appropriate for estimating divergence times. But, extensive analyses of simulated data are required to fully explore the sufficiency of current substitution models and understand whether the difference between simple and complex models can be interpreted as a problem with simple models, which is beyond the scope of this article and an exciting future direction of research.

## Materials and Methods

### Empirical data acquisition

We selected eight large-scale empirical datasets distributed across the tree of life. Species groups, data types, sequence lengths, sequence counts, calibration counts, branch rate model, the number of partitions, and the original substitution models used in the majority of partitions are summarized in **Table 1**. These studies used complex substitution models (e.g., GTR + Γ) along with one or multiple partitions, and none of the studies selected the JC or Poisson models as the best model for any partitions. All empirical data were analyzed initially in MCMCTree (Yang 2007) to estimate the divergence times, except the ‘Spiders’ data that was analyzed in RelTime and then reanalyzed in MCMCTree by Mello et al. (2017). In all analyses, we used the published topologies. We obtained the published divergence times and credibility intervals (CrIs) from the original studies, except for ‘Mammals A’ and ‘Eukaryotes & Prokaryotes’ datasets. Because the original studies of these two datasets did not provide CrIs of Bayesian time estimates, we reproduced timetrees for these two datasets with the same settings as used in the original studies with MCMCTree (v4.9h).

Because analyses of long sequences can require long computational times, mainly in ML branch length calculations, we used the original alignments when they were shorter than 10,000 sites (see **Table 1**). Otherwise, we randomly selected 10,000 sites from the original alignments (10K datasets) for all phylogenetic analyses. The exception was the ‘Plants’ data, for which the original study (Morris et al. 2018) showed that similar time estimates were obtained by using the full alignment (856,439 sites) and a trimmed subsample with high site coverage (2,217 sites). We therefore used the 2,217 sites data for the ‘Plants’ dataset. The subsampled alignments were used in all following analyses of simple and complex models. All empirical datasets are available at https://github.com/cathyqqtao/timing-and-model-complexity.

### Relationship between Bayesian time estimates using simple and complex models

We estimated divergence times in MCMCTree (v4.9h) using simple models with topologies and calibrations from the original studies. All MCMCTree analyses were conducted using the approximate likelihood calculation. The substitution models used in the original studies were employed as the complex models (**Table 1**). Jukes-Cantor (JC) and Poisson models without the assumption of rate variation across sites under the gamma distribution were used as the simple model for nucleotide and amino acid sequences, respectively. For the ‘Plants’ data, we also estimated divergence times using Kimura 2-parameter (K2) (Kimura 1980), Hasegawa-Kishino-Yano (HKY) (Hasegawa et al. 1985) and Tamura-Nei (TN) (Tamura and Nei 1993) models with and without a gamma parameter for accounting for rate variation across sites, and the GTR model without a gamma parameter. We used a single partition in all simple and complex model analyses, although multiple partitions might be used in the original studies. However, we used 29 partitions for the ‘Eukaryotes & Prokaryotes’ data analysis since the original research (Betts et al. 2018) showed a strong influence of partitioning on time estimation. We used the same rate models, prior settings (e.g., tree prior and overall rate prior), and calibration constraints and densities as published in the original studies for all analyses. Two independent runs were conducted to ensure convergence and that ESS values were higher than 200 after removing 10% burn-in samples for each run.

We first examined whether the use of a single partition and subsampled alignments would significantly impact the time estimates generated by the complex model analysis. Therefore, we compared the published times obtained using the full datasets and multiple partitions with times estimated using the subsampled alignments and a single partition. Concordant time estimates were found in all empirical datasets (**Fig. S7**). Thus, we considered the effect of data subsampling and partitioning to be small for the empirical datasets analyzed. Our observations are consistent with studies showing that site subsampling has a limited impact on the accuracy and precision of time estimates (dos Reis and Yang 2013; Zhu et al. 2015). Therefore, we used the divergence times, and CrIs obtained using the subsampled alignments, the original complex substitution model, and a single partition as the inferences from complex models when a reanalysis was needed. We then compared them with the results obtained using the subsampled alignments, simple models, and a single partition to eliminate any site-subsampling bias, and observed good linear relationships (**Fig. 3a**). We also found similar linear trends in direct comparisons between time estimates from simple models and published times for all datasets (results not shown), indicating that the length of sequences has limited impact on the robustness of time estimates to substitution model complexity. We also computed the mean of relative absolute error between times estimated under simple and complex models for each dataset by using 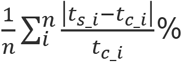, where *t*_c_i_ and *t*_s_i_ is the node time estimated under the complex and simple model for node *i*, respectively, and *n* is the number of nodes. Although we only used MCMCTree for inferring Bayesian divergence times, we expect results to be similar when using other Bayesian dating software, e.g., BEAST2 (Bouckaert et al. 2014), because previous studies have shown that different Bayesian dating software packages tend to generate similar time estimates when priors and calibration constraints are consistent (Warnock et al. 2012; Warnock et al. 2015).

### Influence of calibrations on the relationship of Bayesian times between simple and complex models

We re-estimated divergence times in MCMCTree (v4.9h) using both simple and complex models without any internal calibrations to examine whether the use of many calibrations constrained the final time estimates and concealed bias caused by the use of simple models. In this case, only the age of root was constrained. A single partition and the subsampled alignments were used in the analysis. All other priors, including the root calibration, were the same as those used in the original studies. We also investigated whether the use of root calibration caused a linear relationship between time estimates under simple and complex models. We performed Bayesian analysis for each dataset using a single internal calibration that was randomly selected from all the internal calibrations used in the original study and only the maximum constraint (*t*_max_) for the root. The use of only a maximum constraint for the root results in a diffused uniform density [0, *t*_max_].

### Testing the need for multiple-hits correction in pairwise distance estimation

We tested whether the concordance of times between simple and complex models arose because minimal multiple-hits correction was needed, which was often true for datasets that contained recently-diverged or slow-evolving taxa (Nei and Kumar 2000). We estimated pairwise sequences distances using simple and complex models in MEGA X (Kumar et al. 2018). We used TN and Jones-Taylor-Thornton (Jones et al. 1992) models with a gamma parameter estimated by the ML method as the complex model for nucleotide and amino acid datasets, respectively.

### Testing the relationship of divergence times estimated using simple and complex models for a non-Bayesian approach

We estimated divergence times using the RelTime method (Tamura et al. 2012; Tamura et al. 2018) in MEGA X. A single partition was used in these analyses. No calibrations are required for RelTime analyses. To directly compare the dates estimated by RelTime when no calibrations are used, we normalized time estimates to the sum of node ages. This normalization is simply a *post hoc* scaling of relative times and is not the same as assigning calibrations in the Bayesian approaches in which tree priors interact with calibration probability densities. Outgroups were removed in the time comparisons because RelTime does not produce time estimates for sequences in the outgroup.

### Testing the relationship of branch lengths and node-to-tip distances estimated using simple and complex models

For each dataset, we estimated ML branch lengths using both simple and complex models and the published topology in MEGA X. A single partition was used in all ML analyses. We compared the branch lengths estimated using simple and complex models to obtain the relationship. We then calculated the node-to-tip distances using the resulting ML tree. For each node, the node-to-tip distance is the sum of the lengths of all paths from this node to all descendent tips divided by the total number of descendant tips.

### Testing relationships of branch lengths and node-to-tip distances for different branches and sequence divergences

For each dataset, we compared branch lengths and branch times estimated using simple complex models for short, intermediate, and long branches. Branch length categories were assigned by comparing individual branch length to the mean branch length across a given tree. Long branches were longer than one standard deviation from the mean value of all branches. Short branches were those with lengths shorter than the mean value. The remaining branches were classified as intermediate branches. We also compared node-to-tip distances from simple and complex models for shallow, intermediate, and deep sequences based on the timetree inferred using the complex model and no internal calibrations. The shallow region is the period spanning from 0 Myr to 30% of the root age. The deep region for all datasets, except for the ‘Eukaryotes & Prokaryotes’ data, is the period spanning from 70% of the root age to the root age. For the ‘Eukaryotes & Prokaryotes’ data, the deep region is the period spanning from 50% of the root age to the root age because all internal nodes are younger than 70% of the root age. The remaining timespan belongs to the intermediate region for all datasets. We computed the slopes of node-to-tip distances and of divergence times for nodes that were located in shallow, intermediate, and deep divergences regions.

### Testing the linearity of relationships among substitution model complexity, number of ingroup sequences, and the dispersion of branch lengths

We first randomly sampled ten ingroup sequences from the full ‘Plant’ dataset (99 ingroup + 4 outgroup sequences). We then used “expanded sampling” to generate datasets with 20, 30, 40, 50, 60, 70, 80, 90 ingroup sequences, so that the larger datasets always contained the sequences in the smaller datasets. For example, we kept ten sampled sequences and sampled another ten sequences to generate a dataset with 20 ingroup sequences. We repeated this procedure 20 times, so we had 20 replicates for each number of ingroup sequences. We estimated branch lengths for each replicate using JC, K2, HKY, TN, GTR, and the model used in the original study, GTR + Γ. We compared the branch lengths estimated using the simpler models with those estimated using the GTR + Γ model and computed the coefficient of determination of linear regression through the origin (*R*^2^). Therefore, for each subsampled category, we had 20 *R*^2^ of branch lengths compared between analyses using the GTR + Γ and a corresponding simpler model.

### Testing the relationship of branch lengths and times estimated using the simple model and models that are non-time-reversible and non-stationary

To examine whether the linear relationships of divergence times and branch lengths between simple and complex models are unique phenomena for models in GTR class, we compared the ML branch lengths estimated using the simple model (JC) and models that are non-time-reversible or non-stationary for all nucleotide datasets. We obtained the branch lengths under non-time-reversible (unrestricted) model (model = 10) (Yang 1994) and non-stationary model (nhomo = 3) with a single partition in baseml (v4.9h) (Yang 2007). Because the direct usage of non-time-reversible and non-stationary models is not allowed in MCMCTree for estimating divergence times, we obtained time estimates using the RelTime method with the ML trees produced by baseml and without calibrations. We then normalized time estimates produced by RelTime to the sum of node ages to obtain the relationship.

### Simulation

To assess the impact of sparse taxon sampling and long branches on divergence time estimation between simple and complex models, we conducted a computer simulation. We used a completely unbalanced tree with 16 tips as the model timetree (root age = 3 time units, **Fig. S6a**) and simulated sequences under the strict clock with different mean rates. All sequences were simulated using SeqGen (Grassly et al. 1997) under the GTR + Γ (α = 0.25) model with 5,000 base pairs and a biased base composition (T = 0.25, C = 0.33, A = 0.31, and G = 0.11). We set the mean evolutionary rate of 0.1 substitutions per site per time unit as the baseline case (1×) to make the simulated distribution of pairwise sequence distances (estimated under the TN + Γ model) to be similar to that produced from the analysis of all eight empirical datasets analyzed (1× in **Fig. S6b**). Then we accelerated the mean rate to be 2-, 4-, 6-, 8-, and 10-times faster. For the fastest rate simulated (10×), the median of pairwise sequence distances was 4.7 substitutions per site, which is much larger than the empirical value (**Fig. S6b**).

We inferred divergence times using MCMCTree (v4.9h) using the GTR + Γ and the JC model. Because the likelihood ratio test rejected the assumption of the strict clock (*P* < 10^−20^) when the JC model was used due to uneven underestimation of branch lengths, we relaxed the molecular clock when estimating divergence times. We used three different calibration strategies to investigate the impact of calibration positions on time inference: a precise calibration at a shallow node (o – p divergence in **Fig. S6a**) and a diffused root calibration; a precise calibration at a middle node (j – p divergence) and a diffused root calibration; and a precise root calibration. All priors (e.g., mean evolutionary rate) were set to be as the true values in all Bayesian analyses. Simulated datasets and prior settings are available at https://github.com/cathyqqtao/timing-and-model-complexity.

## Data availability statement

All empirical and simulated datasets are deposited to Github (https://github.com/cathyqqtao/timing-and-model-complexity).

## Acknowledgments

We thank Drs. Sergei Pond, Heather Rowe, Maria Pacheco, Ananias Escalante, Koichiro Tamura, Antonia Chroni, Jeffrey Thorne, Mario dos Reis and two anonymous reviewers for critical comments and editorial suggestions. We also thank Mario dos Reis for suggesting the phylogeny, calibration strategies, and simulation parameters in Fig. S6. This research was supported in part by grants from the National Institutes of Health (NIH GM0126567-02), National Science Foundation (NSF 1661218), and National Aeronautics and Space Administration (NASA NNX16AJ30G) to SK.

