## Supplementary material for "Relative efficiencies of simple and complex substitution models in estimating divergence times in phylogenomics"

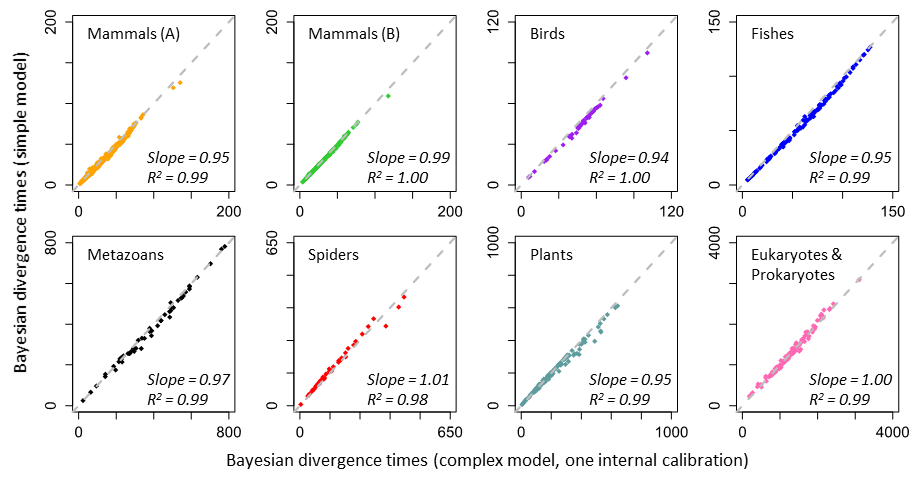

**Figure S1.** Comparison of divergence times obtained using simple and complex models when one internal calibration and a diffused root calibration are used in Bayesian analyses. The time unit is millions of years. The gray dashed line represents the equality of time estimates. The slope and coefficient of determination (*R*^2^) for the linear regression through the origin are shown. References for datasets are listed in figure 2.

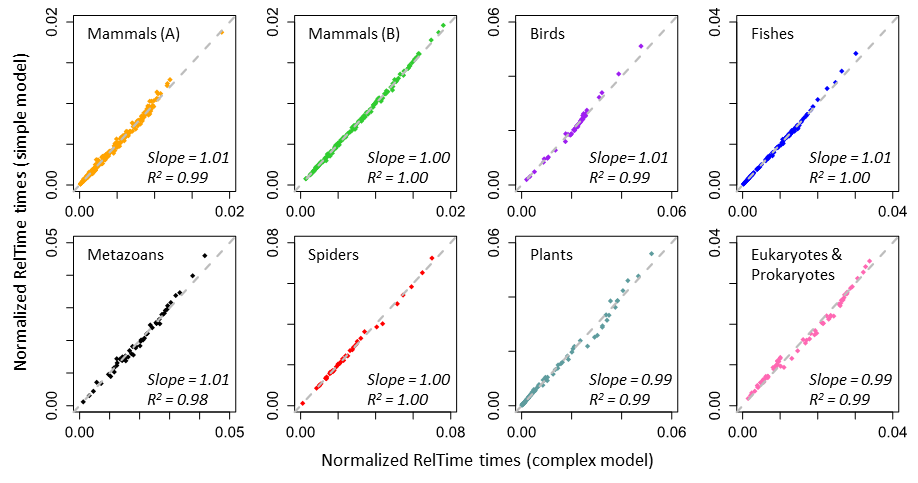

**Figure S2.** Similar time estimates are produced by using simple and complex models without any calibrations in RelTime. Time estimates are normalized to the sum of node ages. The gray dashed line represents the equality of time estimates. The slope and coefficient of determination (*R*^2^) for the linear regression through the origin are shown. References for datasets are listed in figure 2.

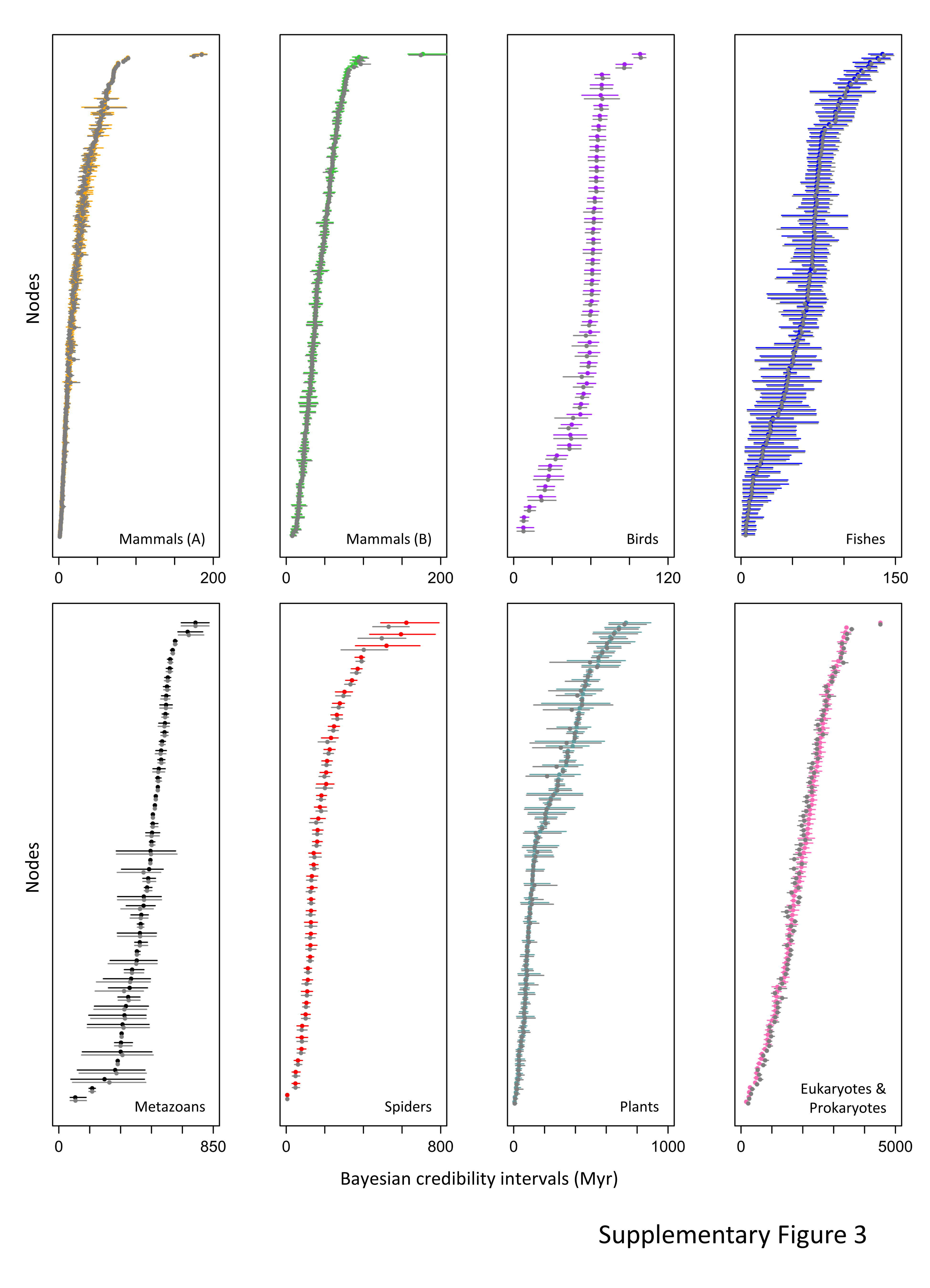

**Figure S3.** Comparisons of Bayesian 95% credibility intervals of divergence times obtained using simple and complex models, when applying all the calibrations. Colored and gray lines represent 95% credibility intervals inferred by using the complex and simple model, respectively. Dots are point estimates of divergence times.

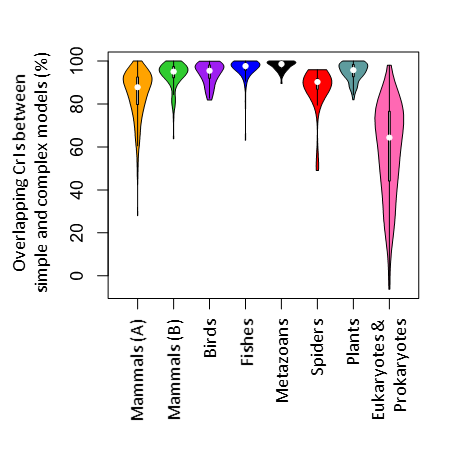

**Figure S4.** Percentage of overlapping regions between Bayesian 95% credibility intervals (CrIs) obtained using simple and complex models, when applying all the calibrations. The percentage of overlapping region for each node is calculated by dividing the amount of overlap of CrIs from simple and complex models by the width of the CrI estimated using the complex model.

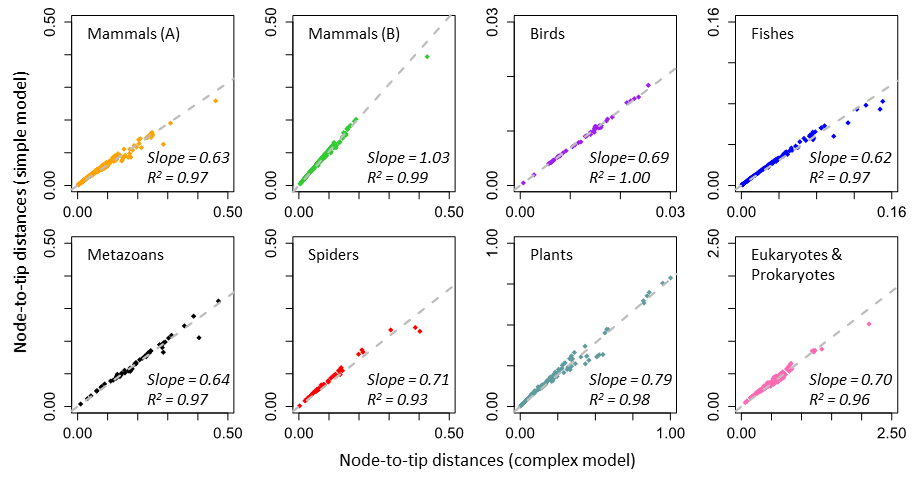

**Figure S5.** Linear relationships between node-to-tip distances obtained by using simple and complex models. The gray dashed line represents the best-fit linear regression through the origin. The slope and coefficient of determination (*R*^2^) for the linear regression are shown.

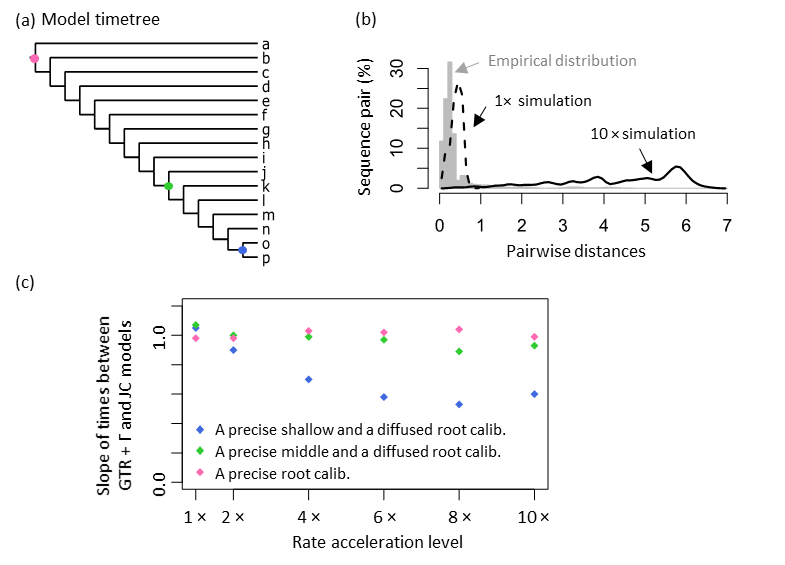

**Figure S6.** (a) A timetree used as the model tree for simulations. Dots represent the position of shallow (blue), middle (green), and root (pink) calibrations. (b) Distribution of pairwise distances for tested empirical datasets (gray bars) and datasets simulated under 1× (dashed black line) and 10× (solid black line) rate scenarios. (c) Slopes of linear regressions (through the origin) between times estimated using the JC and GTR + Γ models for datasets simulated under 1×, 2×, 4×, 6×, 8×, and 10× rate scenarios when a precise shallow calibration and a diffused root calibration (blue), a precise middle calibration and a diffused root calibration (green), and a precise root calibration (pink) is used.

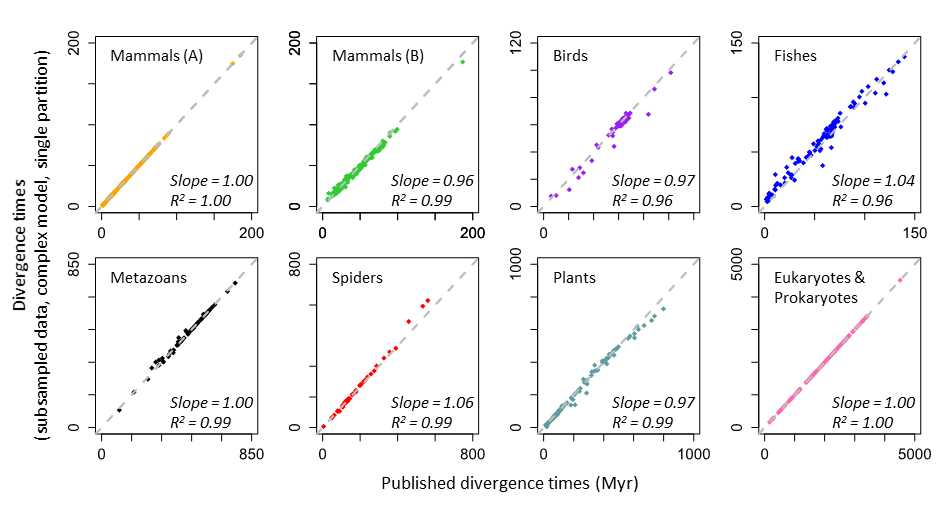

**Figure S7.** Comparison of divergence times obtained using all the data and a small subset of positions in Bayesian analyses. The same complex model, tree topology, and all calibrations are used for each dataset. Gray dashed line represents equality between estimates (1:1 line). The slope and coefficient of determination (*R*^2^) for the linear regression through the origin are shown.
